# General characterization of regeneration in *Aeolosoma viride*

**DOI:** 10.1101/493304

**Authors:** Chiao-Ping Chen, Sheridan Ke-Wing Fok, Yu-Wen Hsieh, Cheng-Yi Chen, Fei-Man Hsu, Jiun-Hong Chen

**Affiliations:** Department of Life Science, National Taiwan University, Taipei, Taiwan; Max Planck Institute of Molecular Cell Biology and Genetics, Dresden, Germany; Stowers Institute for Medical Research, Missouri, America; Graduate School of Frontier Sciences, The University of Tokyo, Chiba, Japan

**Keywords:** *Aeolosoma viride*, Epimorphic regeneration, Cell proliferation, Blastema

## Abstract

Regeneration has long attracted scientists for its potential to restore lost, damaged or aged tissues and organs. A wide range of studies have conducted on different model organisms on both cellular and molecular levels. Current evidences suggest that a variety of regenerative strategies are developed and used by different species, and their regenerative strategies are highly correlated to their reproductive methods. Our present work focused on the freshwater annelid *Aeolosoma viride*, which reproduces by paratonic fission and is capable of complete regeneration. We found out that *A. viride* can regenerate both anterior and posterior end, even with only 3 segments remained. This process is characterized by epimorphosis that involves large amount of cell proliferation which drives the formation of blastema. Cell proliferation and regeneration successful ratio were significantly decreased when treated with microtubule inhibitor taxol or *Avi-tubulin* dsRNA, which confirmed that cell proliferation served as a key event during regeneration. Together, our data described the regenerative processes of *A. viride*, which includes high level of cell proliferation and the formation of blastema. Furthermore, our findings demonstrated *A. viride* as a potential model for the study of regeneration.

## Introduction

An ability to regenerate lost body part is widely, but non-uniformly, distributed in the animal kingdom [1–3]. Both the capacities and mechanisms of regeneration differ between taxa. T. H. Morgan described the processes of regeneration in two generalized terms, namely morphallaxis and epimorphosis [4]. Morphallaxis is characterized by rearrangement of pre-existing cells and the absence of blastema [5]. Early regeneration in hydra is a classic example of morphallaxis as it requires minimum cell proliferation and redistribution of pre-existing tissues. In epimorphosis, on the other hand, undifferentiated cells or progenitor cells aggregate at the wound sites, and these undifferentiated cells proliferate to form a blastema [6, 7]. Therefore, cell proliferation is required for epimorphic regeneration, and application of drugs that perturb cell cycle progression, such as nodcodazole or hydroxyurea, would prevent normal progression of regeneration [8].

Asexual reproduction is often compared to regeneration, since identical individuals arise from fragments of somatic tissue in both processes [9]. Asexual reproduction in annelids can be divided into two major types: architomy and paratomy. Architomy is fission followed by fragmentation. Contrarily, paratomy required individuals to be fully formed or developed prior to fission [10]. In paratomic reproduction, reconstitution or rearrangement of somatic tissue typically occurs in a predictable growth zone. Previous studies demonstrated that such a growth zones can also regenerate lost body parts after injury in certain species reproduce by paratomic fission [9, 11].

Paratomic fission has been described as a common character of annelids, as it is found in taxanomic groups such as Sipuncula, Serpulidae, Aeolosomatidae, and Naididae [12]. Regeneration is also wide-spreading among annelids [13, 14]. Nevertheless, species differs in their regenerative capacities. For example, the naidid *Pristina leidyi* mostly reproduce by paratomic fission and is able to complete both anterior and posterior regeneration [11]. In contrast, the capitellid polychaete *Capitella teleta* mostly reproduce sexually and is only capable to partially regenerate its posterior end [15]. Such a difference may reflect a connection between asexual reproduction by paratomic fission and regeneration [9, 12, 16].

Aeolosomatidae are polychetae that mostly inhabit in fresh water environment [17]. Approximately 30 species have been identified in this family, and predominantly reproduce by agametic reproduction [10, 18]. Most aeolosomatid species reproduce by paratomic fission to create clonal individuals, and an ability to regenerate the anterior and posterior segments after injury has been demonstrated in some species of this family [19]. However, there is no detailed descriptive characterization of the regeneration in this group of annelids. In this study, we provided a detailed description of the regeneration process in *Aeolosoma viride*. It was observed that this worm can restore its lost body parts, both anteriorly and posteriorly, within 3–5 days. Furthermore, we demonstrated that a high level of cell proliferation occurs at the wound site and that such a cell proliferation activity is critical for the completion of regeneration. Using pulse-and-chase experiments, we were able to show that cell migration has no particular role in anterior regeneration of *A. viride*. Finally, our study demonstrated the potential of *A. viride* to be an excellent model for regenerative studies.

## Results

### General and reproductive characteristics of *A. viride*

*Aeolosoma viride* is a 2-3 mm length semi-transparent annelid contained 10 to 12 segments with pairs of chaetae at each side of the worms. The intact worm has a ring-like mouth structure that can be observed at both dorsal and ventral side of the peristomium (1^st^ segment). Mouth is connected to an enlarged digestive tract located at the center of the worm, which can be easily observed under a dissecting microscope. *A. viride* exclusively perform paratonic fission to asexually reproduce one progeny from its posterior end (Fig. 1). The paratonic fission started by extension of the posterior end. Worms exceeding 12 segments with an unusual lengthy posterior region could be easily observed (Fig. 1C). The anterior portion of the offspring gradually developed and formed a head shape tilted apart from the parental worm (Fig. 1D). At this stage, the parent and the progeny can be easily distinguished. Finally, the offspring worm separates apart from the parental worm, and becomes an individual worm (Fig. 1E).

**Figure 1.**
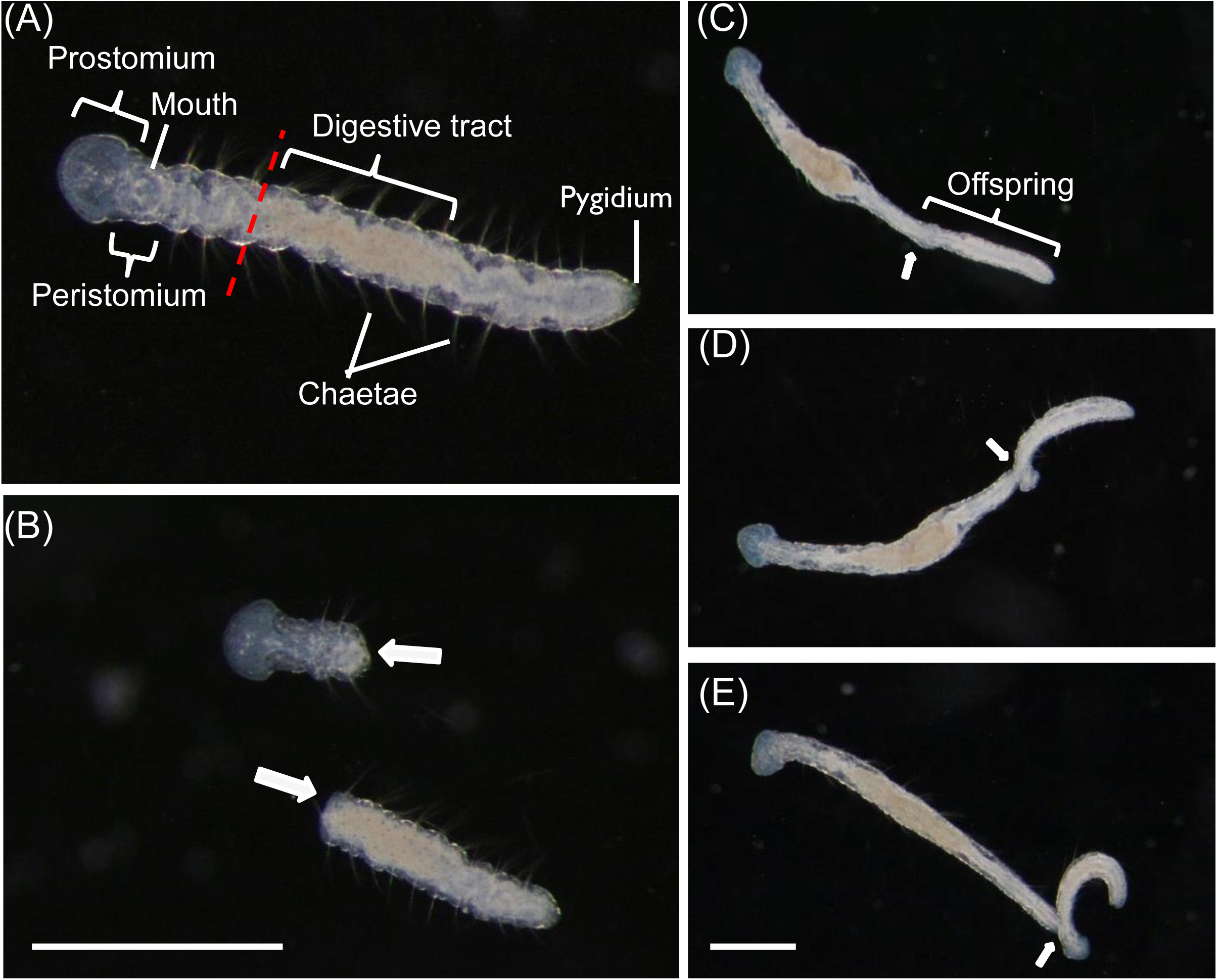
The morphology and paratonic fission of intact *A. viride*. (A) The first segment of intact *A. viride* has a prostomium and a peristomium with a mouth. Average *A. viride* contains 10 to 12 segments with pairs of chaetae. The transparent body has an enlarged digestive tract located at the center of its body. The pygidium is located on the last segment of the posterior end. The red dashed line indicated the amputation site in front of enlarged digestive tract. (B) After the worm was amputated at the site indicated in (A), the two fragments will individually proceed anterior or posterior regeneration (indicated by white arrows). (C-E) The process of paratonic fission separated an intact worm into two individuals. Worms with unusual lengthy posterior region could be easily observed (Fig. 1C). The anterior portion of the offspring gradually developed, and tilted apart from the parental worm (Fig. 1D). The offspring worm break apart from the parental worm, and become an individual worm (Fig. 1E). The white arrow indicated interface between parent and offspring worm. Scale bar: 1 mm.

### Anterior regeneration in *A. viride*

Anterior amputation is performed in front of the enlarged digestive tract. Immediately after amputation, the worm severely twisted; fluids and food residues gushed from the inner cavity. At 3 hours post amputation (hpa), the regenerating *A. viride* adhered on the bottom of culture chamber, and its wounded area remained rough and uneven. The rough wounded area became smoothened within 6 hpa. At 12 hpa, a small amount of tissue became lighter and clearer in color, started to develop at the regenerating area. This tissue will later be proven to be the regenerative blastema. The protruding regenerative blastema expanded from the center of the wounded area and became apparent from 24 to 48 hpa. Vertical contraction could be observed from 72 hpa, but the regenerating worms were still unable to move freely. The reappearance of ring-like structure characterized mouth formation, and a tubular structure (esophagus) extended to connect with the enlarged digestive tract at 96 hpa. After 96 hours of regeneration, the regenerating head area gradually bulged and became wider than other body segments. The prostomium expanded from 96 to 120 hpa (Fig. 2B). During the entire process of regeneration, the regenerating *A. viride* remain adhere on the bottom of culture chamber. Most of them could swim freely around 120 hpa, which was considered as an indicator for successful regeneration in *A. viride*.

**Figure 2.**
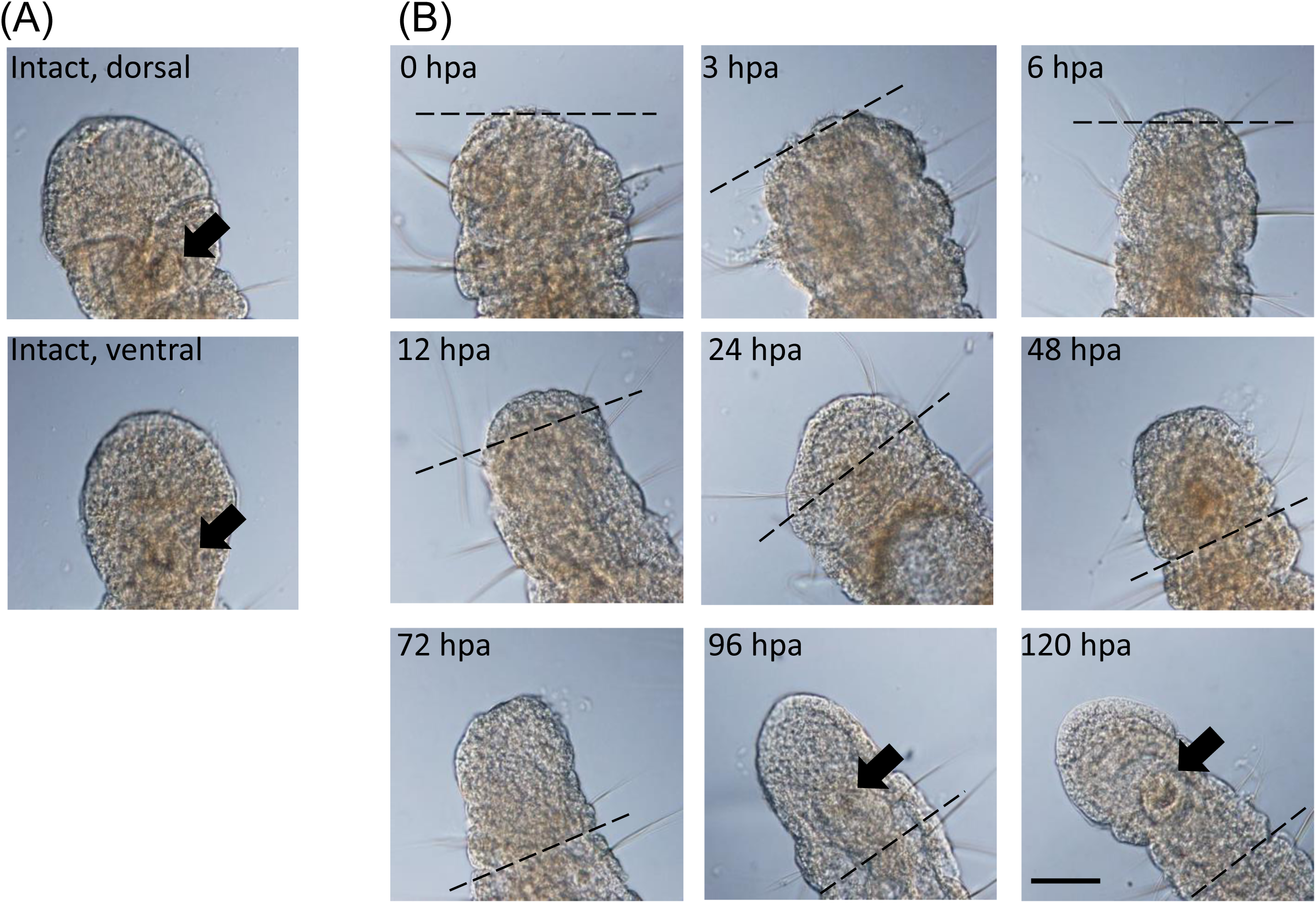
Anterior regeneration in *A. viride*. The worms were amputated in front of the enlarged digestive tract, and the external morphology was observed in intact (A) and 0 to 120 hpa (B) during regeneration. The intact worm has a mouth that can be observed at both dorsal and ventral side of the peristomium at the first segment. During anterior regeneration, the rough wounded area became smooth during wound closure within 3 to 6 hpa. The protruding regenerative blastema became apparent from 24 to 48 hpa in most regenerating *A. viride*. Mouth formation, could be observed around 96 hpa. After 96 hours of regeneration, the cephalic shape started to bulging. Then, the prostomium expanded from 96 to 120 hpa. The amputation site was labeled by black dotted line and the black arrow indicated the re-opening of mouth. Scale bar: 50 μm.

### The regenerative capacity of *A. viride*

For characterization of the regenerative capacity, either the anterior or the posterior ends of worms were amputated. Amputated worms with three segments could still complete regeneration, even the successful ratio was around 40%. Over 60% of worms with 6 or 9 posterior segments completed their anterior regeneration at 5 day post amputation (dpa) (Fig. 3). Posterior regeneration shared a similar pattern. Over 53.3% worms with three anterior segments completed their posterior regeneration. Worms with 6 or 9 anterior segments increased successful ratio of posterior regeneration over 80% at 3 dpa (Fig. 3). Since the anterior segments of *A. viride* contain the mouth and the brain, their complexity and importance outweigh the posterior segments. Therefore, all following experiments in this study were conducted on the anterior end.

**Figure 3.**
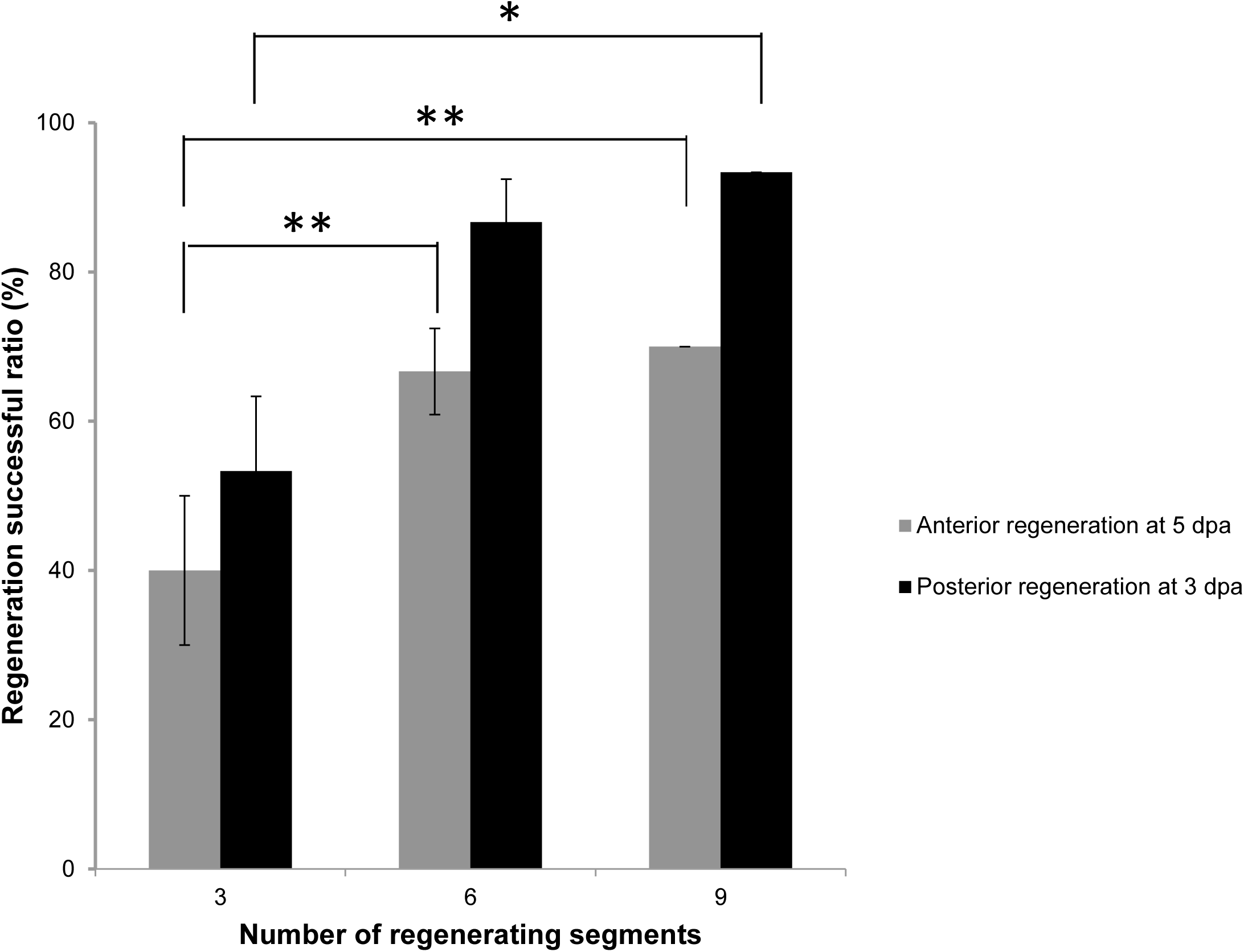
Minimum segments required for successful regeneration in *A. viride*. Worms were amputated into different number of segments. The successful ratio of either anterior or posterior regeneration in amputated worms regenerating from different segments were observed at 5 or 3 dpa. The minimum segments required for successful regeneration in *A. viride* was 3. All data represented the mean ± s.d. from at least three independent duplicate experiments (n = 10). One-way ANOVA was performed to determine the significance of success rates compared to each group. *: *p* <0.05; **: *p* < 0.01.

### Cell proliferation in the regenerative process of *A. viride*

EdU signal showed proliferating cells continuously present in *A. viride*. The worms were also stained with a nuclear dye Hoechst 33342 to conform the EdU labeling (Fig. 4A). Intact and regenerating worms were incubated in EdU solution 12 hours prior to fixation. Normally, the EdU^+^ cells were significally detected at the posterior end of intact worm, the area of asexual reproductive zone, which is the connecting interface between the parent and the future offspring. The EdU signal was also detected at the anterior mouth with minor EdU signals randomly distributed in intact and regenerating worms from 0 to 12 hpa. However, signals posterior to the amputation site quickly diminished after 24 hpa. During regeneration, the EdU signal became concentrated at the regenerating blastema, and the strongest signal was observed at 24 and 48 hpa, then gently decreased from 72 to 120 hpa. At 96 and 120 hpa, EdU^+^ cell formed a ring-like shape at the center of head, which indicated the development of mouth in the regenerating worms (Fig. 4C).

**Figure 4.**
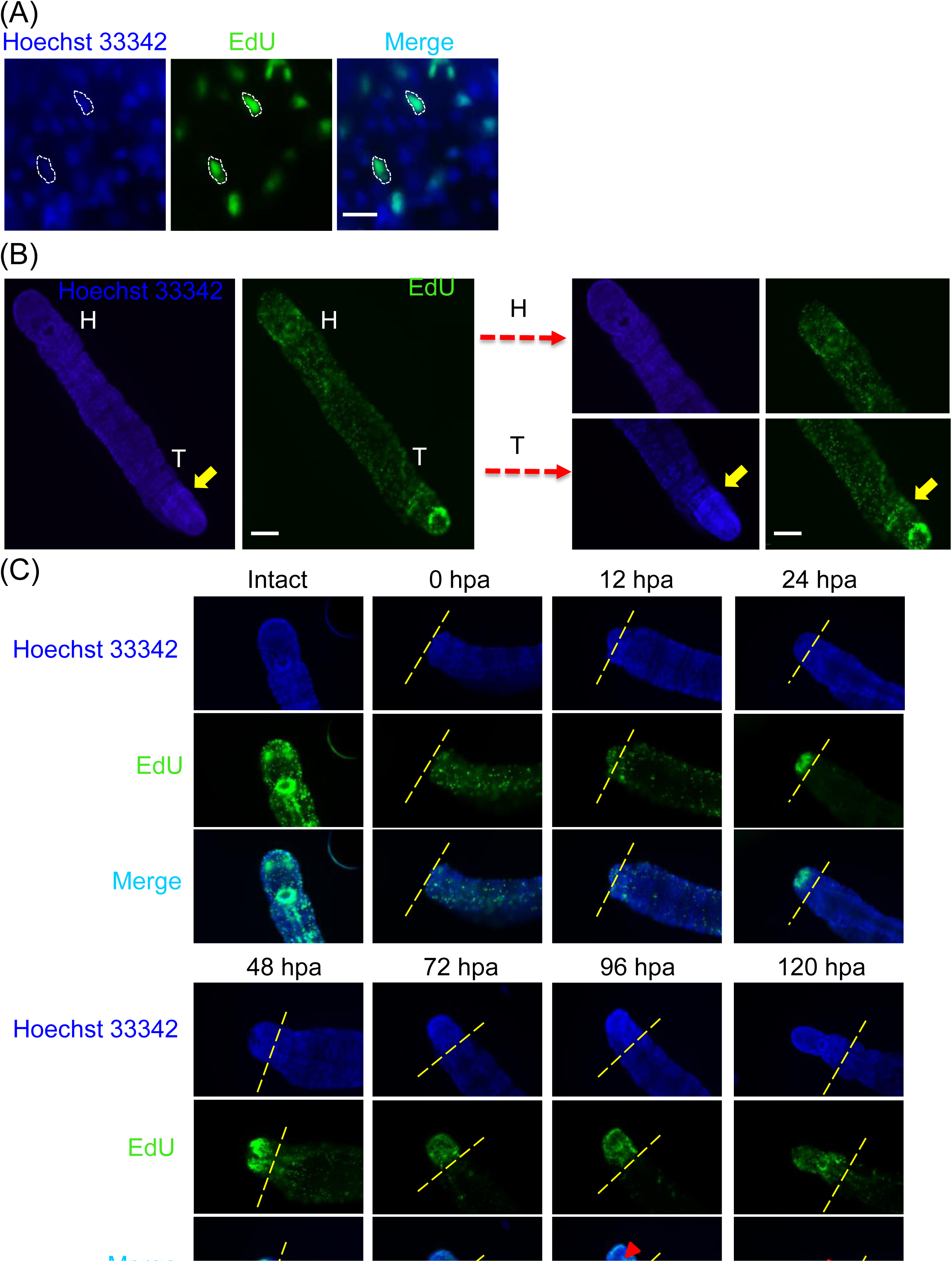
Cell proliferation was detected by EdU labeling during anterior regeneration in *A. viride*. Worms were incubated in 100 μg ml^−1^EdU for 12 hours prior to collecting the specimen at different time points after amputation. (A) EdU signal (green) reveals cell proliferation conformed by nuclei stained with Hoechst 33342 (blue). The white dotted line indicates the colocalization cells in three channels. Scale bar: 10 μm. (B) The worms were labeled with Hoechst 33342 and EdU in intact worm. Figure on the right were magnified pictures of the head or tail regions. The yellow arrow indicates the asexual reproductive zone. Since the asexual reproductive zone undergo continued cell proliferation to produce progeny, the area showed stronger signal of Hoechst 33342 which indicates the presence of larger amount of cells. H: head, T: tail. (C) Cell proliferation was detected on anterior regenerating site at different time points. Prior to amputation, the head, mouth showed the strongest EdU signal. Minor proliferating cells randomly distributed throughout the body. After 12 hpa, EdU signal became focused at the wounded site. Edu signal completely diminished post the amputation site after 24 hpa. The EdU signal peaked at 24 and 48 hpa, then gradually diminished. The amputation site is labeled by yellow dotted line. Red triangle indicated the reappeared mouth. Scale bar: 100 μm.

### Cell proliferation is required for anterior regeneration

*Aeolosoma viride* were treated with taxol, an inhibitor of cell proliferation that works by interfering with the normal function of microtubules. As previous result has shown, the largest amount of proliferating cells was detected at 48 hpa (Fig. 4C). However, the EdU signal in the blastema were apparently reduced by 25 μM of taxol at 48 hpa, which indicated that the proliferating blastema was not properly formed at anterior regenerating site (Fig. 5A). The regeneration successful ratio of *A. viride* treated with 25 μM taxol decreased 80% at 5 dpa and 45% at 7 dpa (Fig. 5B). Also, the bulged head of the regenerating *A. viride* was absent after taxol treatment, and only a tiny blastema was observed at 5 and 7 dpa (Fig. 5C).

**Figure 5.**
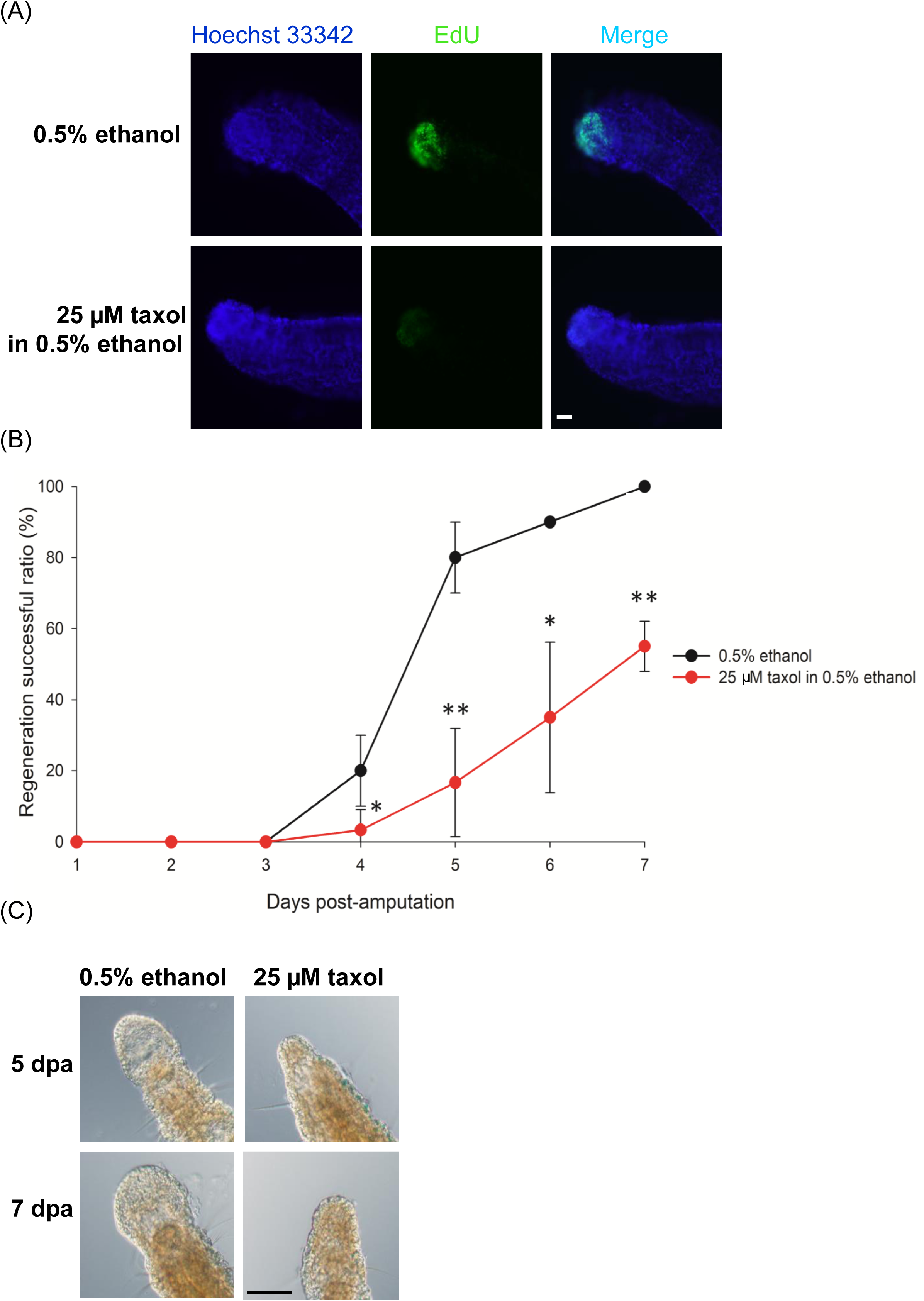
The inhibitory effect of taxol on cell proliferation and anterior regeneration in *A. viride*. (A) *A. viride* weas incubated in EdU 12 hours prior to fixation at 48 hpa. The amount of proliferating cells at the blastema was decreased after 25 μM taxol treatment. (B, C) The regeneration successful ratio was examined from 1 to 7 dpa. Taxol treated worms showed delay and significant decrease compare to the control group at 7 dpa. (D) The head morphology of regenerating worms was obviously affected by taxol treatment. Head formation could be clearly observed in regenerating worm incubated in 0.5% ethanol at 5 dpa, however, only a piece of protruding tissue could be observed in the taxol treated group. At 7 dpa, the ethanol treated group demonstrated normal regeneration, but the taxol treated group barely started the formation of a regenerative blastema. Scale bar: 100 μm. All data represented the mean ± s.d. from at least three independent duplicate experiments. Significant differences relative to control group (0.5% etanol) were denoted by *. *: *p* <0.05; **: *p* < 0.01; ***: *p* < 0.001 using two-tailed unpaired student’s t-test.

To confirm the importance of cell proliferation on regeneration in *A. viride*, we also used gene-specific dsRNA to perform RNA interference. Both feeding and microinjecting dsRNA successfully reduced the expression of *Avi-tubulin* mRNA by 50% compared to *yfp* dsRNA (MOCK) (Fig. 6A). Western blot was conducted to further confirm this result. The predicted molecular weight of tubulin protein is 48.8 kDa. Avi-actin served as loading control, and its predicted molecular weight is 42.1 kDa. The result showed a significant reduction of tubulin protein expression comparing to MOCK RNAi (Fig. 6B). In addition, worms treated with *yfp* dsRNA regenerated normally. But, the regeneration of *Avi-tubulin* RNAi treated animals was significantly inhibited at 5 dpa. The group of feeding dsRNA reduced successful ratio of regeneration by 45%, whereas the group of injecting dsRNA reduced by 35% (Fig. 6C).

**Figure 6.**
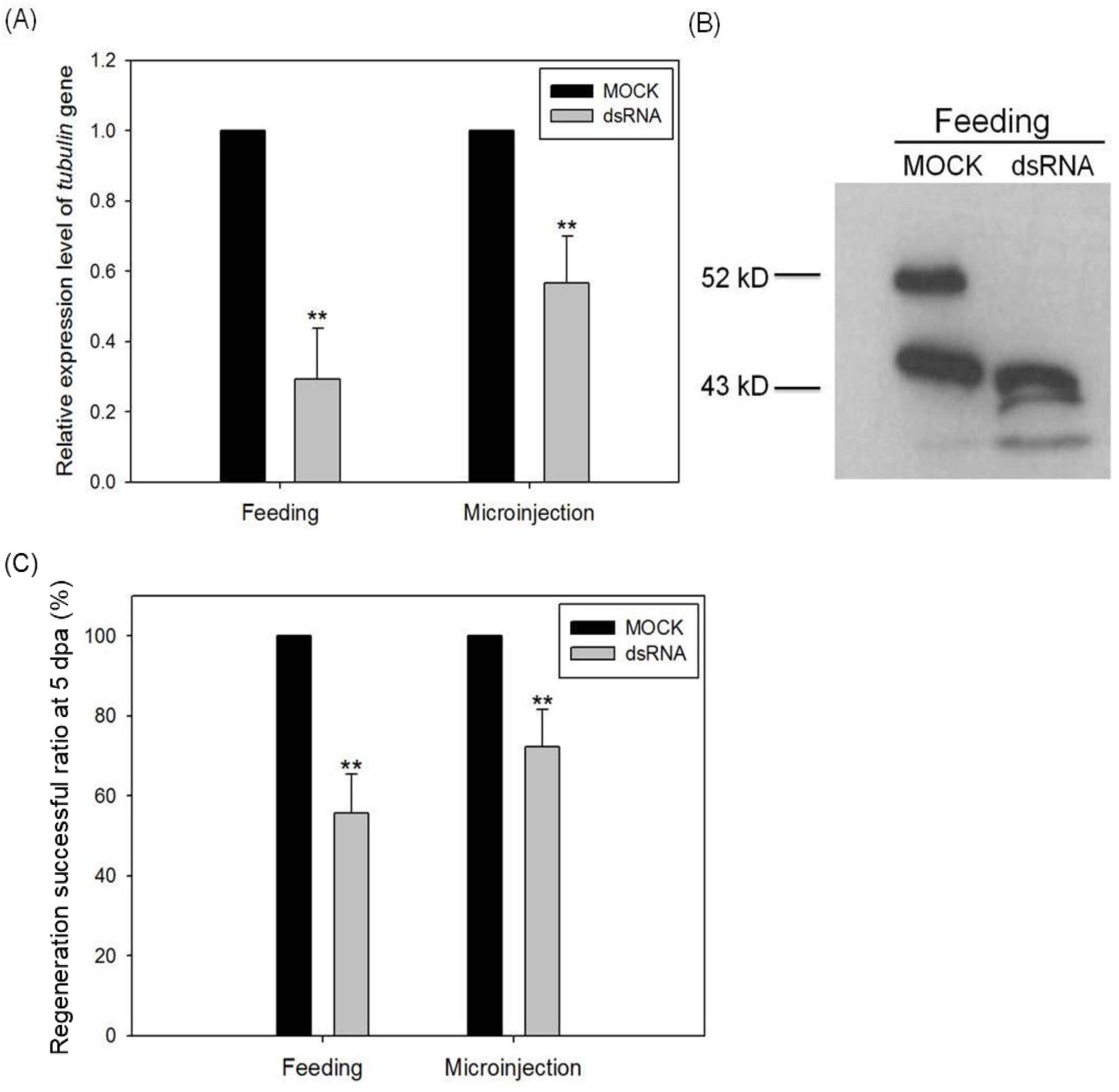
The inhibitory effect of *Avi-tubulin* microRNA on anterior regeneration. Either the expression level of *Avi-tublin* mRNA (A) or protein (B) expression levels o were detected to assay the knock-down efficiency. The inhibitory effect of regeneration successful ratio by *Avi-tubulin* RNAi was observed at 120 hpa (C). One-way ANOVA was performed to determine the significance of mRNA, protein expression or successful ratio compared to control group. **: *p* < 0.01.

### Cell migration in the regeneration of *A. viride*

Some speculated that the source of stem or progenitor cells in the regenerative area could be migration [20]. Stem cell migration not only is fundamental to embryonic development, but also indispensable to adult tissue homoeostasis and repair [21].

To examine the role and importance of cell migration during anterior regeneration of *A. viride*, pulse-chase experiment with EdU labeling was performed. Only a few EdU^+^ cells were detected at the surface of the newly regenerated tissue at 24, 72 and 120 hpa (Fig. 7). However, the distribution of EdU^+^ cells around non-regenerated area was evident from 24 to 120 hpa.

**Figure 7.**
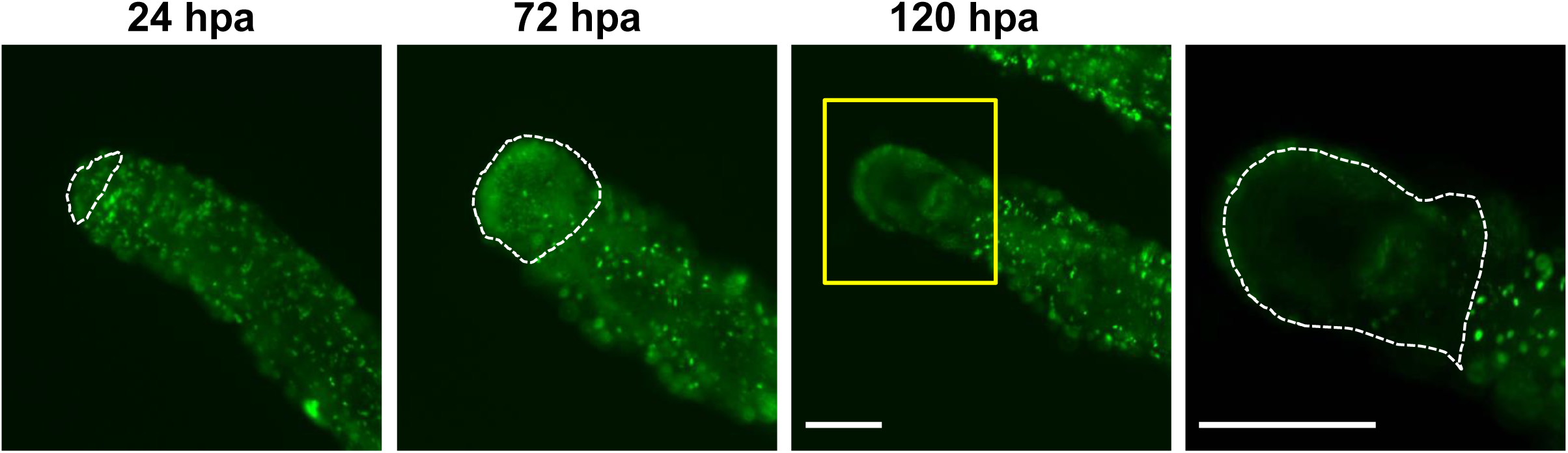
Limited cell migration is observed at anterior regenerating site of *A. viride*. The proliferating cells were incorporated by EdU for 24 hours prior to amputation in intact and regenerating worms. At 24, 72, 120 hours post amputation, minor EdU^+^ cell migrated to regenerating area. Although limited signals could be detected at the surface of regenerating worms, the number of proliferating cells significantly differed from the area behind amputation site. The white dotted line indicates the reappearance of enlarged head. The yellow bracket indicated the enlarged area for the figure on the right side. Scale bar: 100 μm.

## Discussion

*Aeolosoma viride* is the first species in Aeolosomatidae used as a model for detailed regeneration study. In this study, we had carefully evaluated the regenerative capacity of *A. viride*. Regeneration of *A. viride* followed by either anterior and posterior amputation could be completely restored within 5 days and 3 days respectively. Most worms could survive and regenerate with 3 segments. Anterior regeneration showed lower successful ratio than posterior regeneration. It is reasonable to explain that the anterior regeneration is involved the restoration of a more complex structure including head and mouth. Even so, compared with earthworm *Enchytraeus japonensis* that could regenerate into complete individual from 6 fragments [22], and *Eisenia fetida* that could only partially regenerate [23], *A. viride* clearly demonstrated a stronger regenerative ability.

Sexual and asexual reproduction including budding, paratonic fission, and fragmentation have all been documented in annelid reproduction [24]. Previous study has recorded the regenerative capacity and reproductive strategies of many annelids, and discovered that most annelids capable of head regeneration reproduce by asexual reproduction [25]. *Aeolosoma viride* had been recorded that its reproduction is exclusively asexual by paratonic fission [18]. Our study provided detailed imaging on the entire process, and confirmed the idea that animals with asexual reproductive characters typically demonstrated strong regenerative capacity.

Cell proliferation is generally considered to be the key event that occurs at the area of regeneration [26, 27]. The contribution of cell proliferation to regeneration differs across metazoan models [8]. In this study, large amount of EdU^+^ cells present at blastema during regeneration. Combined with the changes in morphology, we observed that anterior amputation in *A. viride* is followed by the formation of blastema with great quantity of cell proliferation during regeneration. Also, significant decrease of proliferating cells was observed at the blastema after treated with mitotic inhibitor taxol. Together, these data demonstrated the importance of cell proliferation in the regenerative process of *A. viride*.

Although variation exists in regeneration ability and mechanism, the widespread phylogenetic distribution of blastema formation in annelid suggested an evolutionary ancient origin of regeneration process. Even though, recent studies indicated that regeneration could not be solely classified as epimorphic or morphollatic, several segmented worm species including *Branchiomma luctuosum* and *C. teleta* have shown limited evidence for morphallaxis [28, 29]. Morphallaxis is characterized by rearrangement of pre-existing tissues, which strongly depend on stem cells or remaining undifferentiated cells migrating to the injured site [30]. Our data supported the hypothesis that *A. viride* carried out anterior regeneration by epimorphosis which is consistent with most other annelids [29, 31–34]. Recent findings inferred that regeneration in annelids involves both cell proliferation and cell migration [35]. And the cellular source of blastema originated from the migration of stem cells [20]. According to our pulse-chase experiment, EdU+ cells were limited in the anterior regenerating site, which means that only a small amount of proliferating cells migrated into the wounded area during anterior regeneration. This allowed us to hypothesize that cell migration served no specific or limited role during anterior regeneration in *A. viride*. There was no evident morphallaxis occurred, however, further experiment with stem cell markers and live-imaging are needed for a definite conclusion.

Limited regeneration studies conducted on annelids model organisms. Hydra and planarian are both considered as the master of regeneration, and often used as the model organisms for regeneration studies [7, 36–39]. But, there are specific restrictions on regeneration studies regarding these commonly used animals. Both planarian and hydra lack sophisticated organ systems and a complex central nervous system, which is far different from vertebrate animals. Hydra regeneration is also classified as morphallatic, which means that they could regenerate under limited cell proliferation [40, 41]. And based on current studies, none of the known vertebrate could regenerate mainly through morphallaxis. Epimorphosis is the major type of regeneration observed in vertebrate animals. Salamander, *Xenopus* and zebrafish are all prominent examples of animal models used in the study of epimorphic regeneration. However, regeneration in these animals is limited to specific organs or tissues. Some might even have restriction on early stage of life [42–44]. Certain species of echinoderms could completely regenerate from a piece of tissues [45], but echinoderm shared very different characteristic from vertebrate animals, such as body symmetry, lack of head, digestive, and circulatory system. Due to the limitations that current models retained, we believe that annelids can serve as different model organism for the study of regeneration. Besides an evolutionary advantage, *A. viride* also possess similar level of experimental advantages compare to many current model organisms. Small molecular inhibitors, molecular tools were both applicable on this worm. First, *A. viride* is easy to maintain and raise in laboratory. Second, *A. viride* has small and transparent body which make them easy to be manipulated and observed during the experimental processes. Last but not least, RNA interference was used in the study of molecular biology to knock down or silence specific mRNA expression. Feeding method commonly designed for *C. elegans* and microinjecting method designed for animal embryo and adult were both functional in this newly introduced model animal [46]. In addition to its regeneration capacity, this small annelid has relative short lifespan and reproductive cycle which is suited for study on aging [47–49].

In conclusion, we examined the regenerative capacity in *A. viride*, especially on anterior regeneration. In addition to strong regeneration ability, *A. viride* has several characters that may make them valuable as a potential animal model for regeneration research.

## Materials and Methods

### Animals

*Aeolosoma viride* was cultured in artificial spring water (ASW, 48 mg L^−1^ NaHCO_3_, 24 mg L^−1^ CaSO_4_•2H_2_O, 30 mg L^−1^ MgSO_4_•7H_2_O, and 2 mg L^−1^ KCl in distilled water, pH = 7.4) at 25°C under 12 hours of day-night cycles. Grounded oat meal serves as major food source. Approximately 500 ± 200 worms were fed with 20 mg powdered oats 3 to 5 times per week. Prior to all experiment, worms were starved in ASW overnight.

### Regeneration experiment

*Aeolosoma viride* without offspring was selected for the following experiment. To analyze the capacity of regeneration, amputation was carried out at the desired segment in *A. viride* (Fig. 1A). The detail morphological changes of regenerating worms were observed with an Olympus DP80 microscope for the following 3 to 168 hours. The successful ratio of anterior or posterior regeneration was determined from morphology and behavior with a dissecting microscope (WILD M8, Leica) at 1 to 7 days post-amputation (dpa).

In the inhibitory experiment, all procedures were similar as described earlier, but the animal were transferred into fresh ASW containing 25 μM taxol (Sigma-Aldrich)

### EdU labeling

Worms were treated with EdU (100 μg ml^−1^) for 12 or 24 hours prior to amputation; 12 to 120 hours prior to fixation. The animals were fixed with 4% paraformaldehyde (PFA) at 4□ overnight. Whole-mount immunohistochemistry detection was processed using the Click-iT EdU Alexa Fluor 488 Imaging Kit (Invitrogen) according to the manufacturer’s protocol.

### RNA interference (RNAi)

*Avi-tubulin* was cloned by our lab (NCBI # AQV09899.1). The RNA interference protocol was modified as described previously [50]. Partial sequence of yellow fluorescent protein (*yfp*, as MOCK group) or *Avi-tubulin* were constructed with L4440 vector and transformed into an RNase III deficient strain competent cell HT115 (DE3) (Yeastern Biotech). *A. viride* was fed with bacteria containing dsRNA for 3 days.

Microinjecting method was modified based on previous study [51]. The *yfp* or *Avi-tubulin* dsRNA were *in vitro* transcribed with T7 polymerase (Ambion). Each individual animal was injected 50 ng of dsRNA through microinjector (Promega) for 3 consecutive days. Worms were then collected for RNA extraction, protein extraction or regeneration study afterward.

### RNA exaction and quantitative real-time RT-PCR (qRT-PCR)

The total RNAs were extracted from 20 worms by using TRIzol and thenreverse-transcribed to cDNA by using SuperScript III Kit (Invitrogen). Transcriptional levels were determined by Bio-Rad iCycler™ (Bio-Rad) using SYBR green system. The primers used to amplify *Avi-tubulin* were 5’-GGTAACAACTGGGCTAAGGG-3’ and 5’-GCGAAGCCAGGCATGAAGAA-3’. *Avi-actin* was used as internal control with specific primers: 5’-ATGGAGAAGATCTGGCATCA-3’ and 5’-GGAGTACTTGCGCTCAGGTG-3’ designed from *Avi-tublin* (NCBI # AQV09898.1). Relative quantification of gene expression was calculated by using the ΔΔCt method. Three technical replicates were used in each real-time PCR reaction, and a no-template blank was served as negative control.

### Western blotting

Animals were homogenized in RIPA buffer (50 mM Tris-HCl (pH 7.0), 150 mM NaCl, 1% Triton X-100, 0.5% C_24_H_40_O_4_, 0.1% SDS with protease inhibitor cocktail (Sigma-Aldrich) and DNase (Promega)). The homogenized samples were mixed with sample buffer (60 mM Tris-HCl (pH 6.8), 25% glycerol, 2% SDS, 14.4 mM 2-mercaptoethanol and 0.1% bromophenol blue), separated by 7.5% polyacrylamide gel, and transferred to a PVDF membrane (Millipore). After blocking with 5% BSA at 25□ for 1 h, the PVDF membrane was incubated with either anti-acetylated α-tubulin (1:8000, GeneTex) or anti-α-actin (1:8000, Santa Cruz) antibodies in the blocking solution at 25□ for 1 h, rinsed three times with PBST, and then incubated with either HRP-conjugated goat anti-mouse IgG (1:3000, Chemicon-Millipore) for α-tubulin or HRP-conjugated goat anti-rabbit IgG (1:8000, Chemicon-Millipore) for α-actin. Finally, patterens were detected on the PVDF membrane by ECL (Bioman).

### Statistics

Data were test for significance using one-way analysis of variance with a Scheffe’s method post *hoc* test or two-tailed unpaired student’s t-test. Probability values of *p* < 0.05 were regarded as statistically significant.

## Acknowledgments

We thank Dr. Kuo Dian-Han and Dr. Lai Yi-Te, who contributed greatly by proving continual guidance, editing and reviewing this manuscript.

## Funding

This study was supported in part by a grant from the Ministry of Science and Technology (MOST, Taiwan, grant no. 103-2311-B-002-017-MY3)

## Availability of data and materials

The datasets generated and/or analysed during the current study are available from the corresponding author on reasonable request.

## Author contributions

All authors designed the experiments. CC, SF and YH carried out the experiments. CC and SF interpreted the data. CC and SF wrote the manuscript. JC reviewed this manuscript.

## Competing interests

The authors declare no competing interests.

